# Long-read genome assemblies for the study of chromosome expansion: *Drosophila kikkawai*, *Drosophila takahashii*, *Drosophila bipectinata*, and *Drosophila ananassae*

**DOI:** 10.1101/2023.05.22.541758

**Authors:** Wilson Leung, Nicole Torosin, Weihuan Cao, Laura K Reed, Cindy Arrigo, C R Sarah Elgin, Christopher E Ellison

## Abstract

Flow cytometry estimates of genome sizes among species of *Drosophila* show a 3-fold variation, ranging from ∼127 Mb in *Drosophila mercatorum* to ∼400 Mb in *Drosophila cyrtoloma*. However, the assembled portion of the Muller F Element (orthologous to the fourth chromosome in *Drosophila melanogaster*) shows a nearly 14-fold variation in size, ranging from ∼1.3 Mb to > 18 Mb. Here, we present chromosome-level long read genome assemblies for four *Drosophila* species with expanded F Elements ranging in size from 2.3 Mb to 20.5 Mb. Each Muller Element is present as a single scaffold in each assembly. These assemblies will enable new insights into the evolutionary causes and consequences of chromosome size expansion.

## Introduction

Genome size spans several orders of magnitude across eukaryotes: some fungi have genomes less than 10 Mb in size while some plant and protozoan species have genomes larger than 100 Gb (Gregory *et al*. 2007). This size variation is not associated with organismal complexity nor with the number of protein coding genes, i.e. “the C-value paradox” (Elliott and Gregory 2015). While it is now clear that variation in the abundance of non-coding (usually repetitive) DNA is the major determinant of genome size variation in eukaryotes, many questions remain, including the evolutionary causes and consequences of the proliferation of non-coding DNA (Gregory 2005).

*Drosophila* genomes have six chromosome arms, known as Muller Elements A – F, arranged in four to six chromosomes (Muller 1940). The Muller F Element is the ancestral X chromosome across all Dipteran species, however prior to the most recent common ancestor of *Drosophila*, the F Element became an autosome and underwent a large reduction in size (Vicoso and Bachtrog 2013). The *D. melanogaster* Muller F Element exhibits characteristics distinct from the other Muller Elements in several ways, including a low rate of recombination, late replication, enrichment of the histone modification H3K9me2/3, and high levels of Heterochromatin Protein 1a (HP1a) and Painting of fourth (Pof) (Larsson *et al*. 2004). The *D. melanogaster* F Element is thus considered almost entirely heterochromatic, with an approximate size of 5.2 Mb (Locke and McDermid 1993). However, the distal 1.3 Mb contains approximately 80 protein-coding genes that show a range of expression levels similar to genes located in the euchromatic regions of the other autosomes (Riddle and Elgin 2018).

Across the *Drosophila* genus, which is paraphyletic (Finet *et al*. 2021; Suvorov *et al*. 2022), overall genome size is fairly constrained. Flow cytometry suggests the largest genome size (∼400 Mb, *D. cyrtoloma*) is only ∼3-fold larger than the smallest (∼127 Mb, *D. mercatorum*) (Bosco *et al*. 2007, Craddock *et al*. 2016, Gregory 2023). However, as inferred from current genome assemblies, the portion of the *Drosophila* Muller F Element containing protein-coding genes shows a remarkable, nearly 14-fold variation in size, ranging from ∼1.3 Mb in the *D. melanogaster* Release 6 assembly to ∼17.8 Mb in the *D. ananassae* dana_caf1 genome assembly (*Drosophila* 12 Genomes Consortium *et al*. 2007; Schaeffer *et al*. 2008).

Prior work in *D. ananassae* found that proliferation of retrotransposons played a major role in size expansion of the F Element (Leung *et al*. 2017). Furthermore, while *D. ananassae* and *D. melanogaster* F Element genes share distinct characteristics — including larger coding spans and introns, greater number of coding exons, and lower codon usage bias - these features are exaggerated in *D. ananassae*, potentially due to the F Element size expansion (Leung *et al*. 2017). However, these observations were based on the analysis of a small portion of the *D. ananassae* F Element (i.e., ∼1.4 Mb from two scaffolds).

Here we have used long-read sequencing and Hi-C scaffolding to construct chromosome-level genome assemblies for three *Drosophila* species that show different degrees of F Element size expansion: *D. takahashii*, *D. kikkawai*, and *D. bipectinata* (Figure 1). We also perform Hi-C scaffolding for a recently generated long-read *D. ananassae* genome assembly (Tvedte *et al*. 2022). All Muller Elements, including the F Element, are present as single scaffolds in each of these assemblies; thus providing a valuable resource for future studies investigating the causes and consequences of size expansion in this chromosome.

**Figure 1.**
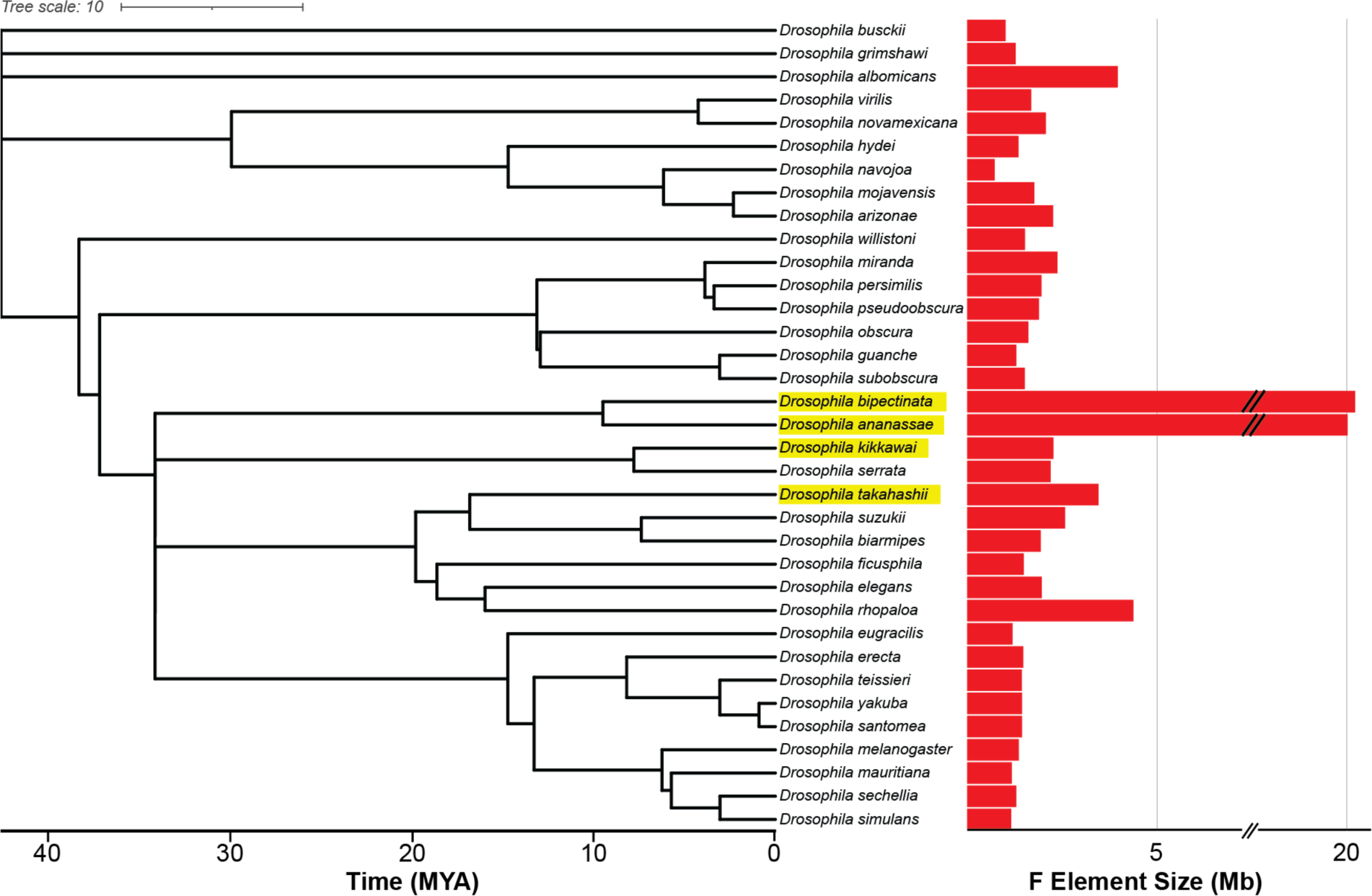
Size variation of the Muller F Element across the *Drosophila* genus. The size of the F Element for 35 *Drosophila* species was estimated based on their NCBI RefSeq genome assemblies (see Methods). The species considered here are highlighted in yellow. Tree topology and timescale were obtained from the TimeTree of Life resource (Kumar *et al*. 2022). Note that the F Element size x-axis is discontinuous due to space constraints.

## Methods & Materials

### Software versions

See Table S1 for the version information and the references for the bioinformatics tools used in this study.

### Muller F size estimation

The F Element sizes in each RefSeq genome assembly were estimated based on the total size of the scaffolds that contain alignments to *D. melanogaster* F Element proteins, transcripts, and coding exons. The RefSeq genome assemblies for 35 *Drosophila* species were obtained from NCBI (O’Leary *et al*. 2016). The *D. melanogaster* proteins, transcripts, and coding exons sequences were obtained from FlyBase (FlyBase release FB2022_03; *D. melanogaster* annotation release 6.46) (Gramates *et al*. 2022). In addition, the *D. melanogaster* release 6 genome assembly was aligned against each RefSeq genome assembly to facilitate the identification of *D. melanogaster* F Element genes that might have moved to another Muller Element in other *Drosophila* species. The proteins, transcript, coding exons, and whole genome alignments are available as evidence tracks (i.e., *D. melanogaster* Proteins, *D. melanogaster* Transcripts, CDS Mapping, *D. melanogaster* Net) on the Genomics Education Partnership mirror of the UCSC Genome Browser (https://gander.wustl.edu).

The *D. melanogaster* proteins were aligned against each RefSeq genome assembly with SPALN (Iwata and Gotoh 2012) with cross-species alignment parameters optimized for *D. melanogaster*: -yX1 -TInsectDm -LS -ya0 -yS100.

*D. melanogaster* transcripts were aligned against each RefSeq genome assembly using BLAT with the parameters -q=rnax -t=dnax -mask=lower. The transcript alignments were analyzed and filtered by utilities provided by the UCSC Genome Browser (Nassar *et al*. 2023). Transcript alignments were analyzed by pslReps to identify the genomic region with the best alignment having a minimum alignment ratio of 0.25 (-minAli=0.25). The alignments were then filtered by pslCDnaFilter using the following parameters: -minId=0.35 -minCover=0.15 - localNearBest=0.010 -minQSize=20 -ignoreIntrons -repsAsMatch -ignoreNs -bestOverlap.

*D. melanogaster* coding exons were aligned against each RefSeq genome assembly with the tblastn program provided by WU-BLAST with the following parameters: -e=1e-2 -topComboN=1 -links -hspsepSmax=10000 -hspsepQmax=1000 -matrix=PAM40 -Q=7 -R=2.

The *D. melanogaster* whole genome assembly was obtained from FlyBase, and aligned against each RefSeq genome assembly using LAST (Kiełbasa *et al*. 2011). The whole genome alignments were then processed using the chain and net alignment protocol and utilities (e.g., axtSort, axtChain, chainAntiRepeat, chainFilter, chainPreNet, chainNet, netSyntenic, netClass, netFilter) developed by the UCSC Bioinformatics Group (Kent *et al*. 2003).

### Strain information

#### *Drosophila kikkawai* strain 14028-0561.14

The original strain of *D. kikkawai* was obtained from the National *Drosophila* Species Stock Center (NDSSC) at Cornell University, and then inbred (11 generations of full-sib crosses) by Professor Artyom Kopp at University of California, Davis (UC Davis). Details regarding the BioSample used for the PacBio whole genome sequencing are available under the accession number <u>SAMN33872896</u>, and the details for the BioSample used to generate the Hi-C data are available under the accession number <u>SAMN34351228</u>.

#### Drosophila takahashii strain IR98-3 E-12201

The original strain of *D. takahashii* was obtained from EHIME-Fly, and then inbred (10 generations of full-sib crosses) by Professor Artyom Kopp at UC Davis. Details regarding the BioSample used for the PacBio whole genome sequencing are available under the accession number <u>SAMN33872897</u>, and the details for the BioSample used to generate the Hi-C data are available under the accession number <u>SAMN34351229.</u>

#### Drosophila bipectinata strain 14024-0381.07

The strain of *D. bipectinata* was obtained from the NDSSC, and then kept by the Elgin Lab and Kopp Lab, but not inbred. Details regarding the BioSample used for the PacBio whole genome sequencing are available under the accession number <u>SAMN33872898</u>, and the details for the BioSample used to generate the Hi-C data are available under the accession number <u>SAMN34351230.</u>

#### Drosophila ananassae strain 14024-0371.13

This strain of *D. ananassae* was originally kept by the *Drosophila* Species Stock Center (DSSC) at the University of California, San Diego, and sequenced by the *Drosophila* 12 Genomes Consortium (*Drosophila* 12 Genomes Consortium *et al*. 2007). The strain of *D. ananassae* used to generate the whole genome assembly produced by Professor Julie C. Dunning Hotopp at the University of Maryland (Tvedte *et al*. 2021) was treated with tetracycline and is cured of *Wolbachia* (see BioSample <u>SAMN13901672</u> for details). The strain of *D. ananassae* used to generate the Hi-C data was maintained independently by the Elgin Lab and was not treated with tetracycline (see BioSample <u>SAMN26507075</u> for details).

In addition to EHIME-Fly and the NDSSC, the strains of *D. kikkawai*, *D. takahashii*, *D. bipectinata*, and *D. ananassae* used in this study are currently available through the Ellison Lab (Rutgers University).

### PacBio long-read sequencing

High molecular weight (HMW) DNA from *D. kikkawai* (adult females), *D. takahashii* (adult females), and *D. bipectinata* (adult males and females) flies was provided by Dr. Bernard Kim at Stanford University. An overview of the DNA extraction protocol has previously been described (Kim *et al*. 2021), and the detailed protocol is available at protocols.io (Y Kim *et al*. 2020).

The library preparation and PacBio sequencing were performed by the McDonnell Genome Institute (MGI) at Washington University in St. Louis. The SMRTbell Express Template Prep Kit 2.0, Sequel Binding Kit 3.0, and Sequel Sequencing Kit 3.0 were used to prepare the samples for single molecule real-time (SMRT) sequencing using the PacBio Sequel system. Each species was sequenced using one 1M SMRT cell in the continuous long-read sequencing (CLR) mode with the 6.0.0.45111 chemistry, and a movie length of 600 minutes. The sequencing data were processed by version 7.0.1.66975 of PacBio SMRT Link.

To assess the quality of the PacBio sequencing data, the Quality Control (QC) tool in SequelTools (Hufnagel *et al*. 2020) and SEQUELstats were used to analyze the subreads and scraps BAM files for each species.

### Assembly of PacBio reads

The PacBio subreads were analyzed by the icecreamfinder.sh script in BBMap to remove adapter sequences and filter potential chimeric reads. Trimmed PacBio subreads that passed the filter and have a read length of at least 5000 nt were used as input to the Canu assembler (Koren *et al*. 2017) with the genomeSize parameter set to 205m. The Canu assemblies then underwent two rounds of polishing by GCpp (with the Arrow algorithm) using the PacBio subreads. The assemblies from the second round of GCpp were then polished using Illumina genomic reads from each species by POLCA [part of the MaSuRCA assembler; (Zimin and Salzberg 2020)] and NextPolish (Hu *et al*. 2020). The Illumina genomic reads used for polishing were obtained from the NCBI Sequence Read Archive (SRA) under the accession numbers <u>SRR345537</u> (*D. kikkawai*), <u>SRR13070706</u> (*D. takahashii*), and <u>SRR6425989</u> (*D. bipectinata*).

### Assembly of Nanopore reads

Nanopore reads were obtained from the NCBI SRA under the accession numbers <u>SRR13070622</u> (*D. kikkawai*), <u>SRR13070623</u> (*D. takahashii*), and <u>SRR13070724</u> (*D. bipectinata*). Adapter sequences and chimeric reads were identified and removed by Porechop. The trimmed Nanopore reads that passed the default Porechop filters were used as input to the Flye assembler (Kolmogorov *et al*. 2019) with the genome-size parameter set to 205m. The Nanopore Flye assemblies then underwent two rounds of polishing with GCpp using the PacBio subreads from the corresponding species. The assemblies from the second round of GCpp were polished by POLCA and NextPolish using the same set of Illumina reads used to polish the PacBio assemblies.

### Assembly merging

For each species, two rounds of quickmerge (Chakraborty *et al*. 2016) were used to combine the PacBio assembly produced by Canu with the Nanopore assembly produced by Flye. In the first round of quickmerge, the Nanopore assembly produced by Flye was used as the query and the PacBio assembly produced by Canu was used as the reference to produce the merged assembly. In the second round of quickmerge, the PacBio assembly produced by Canu was used as the query and the merged assembly produced by the first round of quickmerge was used as the reference. The assemblies produced by the second round of quickmerge were then polished by Hapo-G (Aury and Istace 2021) using the genomic Illumina reads that had been used for polishing the PacBio Canu and Nanopore Flye assemblies.

The polished quickmerge assemblies were analyzed by Purge Haplotigs (Roach *et al*. 2018) to identify haplotigs associated with the primary contigs. Different -low, -mid, and -high parameters were used with the cov command to identify haplotigs based on alignment coverage: *D. kikkawai* (-low 5 -mid 20 -high 95); *D. takahashii* (-low 5 -mid 100 -high 190); *D. bipectinata* (-low 5 -mid 95 -high 190).

### Hi-C scaffolding

All flies were maintained in population cages on molasses agar with yeast paste (https://bdsc.indiana.edu/information/recipes/hardagar.html). Embryos (8 – 16 hours) for each species were collected and dechorionated in 50% commercial bleach for 2.5 minutes at room temperature. Nuclei were isolated from 250 – 500 mg of embryos and fixed in 1.8% formaldehyde for 15 minutes according to a previously published protocol (Sandmann *et al*. 2006). Hi-C libraries were constructed for each species using the *in situ* DNase Hi-C protocol in (Ramani *et al*. 2016) and 150 bp paired end reads were sequenced on an Illumina HiSeq machine. Approximately 116 – 177 million read pairs were generated for each species.

Hi-C scaffolding was performed using the 3D-DNA pipeline (Dudchenko *et al*. 2017) with the following parameters: EDITOR_REPEAT_COVERAGE: 6, SPLITTER_COARSE_STRINGENCY: 70, SPLITTER_SATURATION_CENTILE: 7, SPLITTER_COARSE_RESOLUTION: 50000. A gap size of 500 bp (--gap_size 500) was added between adjacent contigs produced by quickmerge to construct the sequences in the Hi-C scaffolded genome assemblies. Contact maps for each chromosome arm were generated using the *hicPlotMatrix* utility from HiCExplorer (Ramírez *et al*. 2018) with 40 Kb bins.

The snail plots used to assess the quality of the Hi-C scaffolded genome assemblies were generated by BlobTools2 (Challis *et al*. 2020).

### Assembly level classification

The NCBI Assembly Data Model defines a chromosome-level assembly as follows: “There is sequence for one or more chromosomes. This could be a completely sequenced chromosome without gaps or a chromosome containing scaffolds or contigs with gaps between them. There may also be unplaced or unlocalized scaffolds” (https://www.ncbi.nlm.nih.gov/assembly/help/#level). We therefore describe our assemblies as chromosome-level despite the fact that they contain gaps as well as unplaced scaffolds.

### Annotation

The diptera_odb10 (release date 2020-08-05) lineage dataset was used with BUSCO (Manni *et al*. 2021) in “genome” mode to assess the quality of the assembled genomes. The generate_plot.py script provided by BUSCO is used to produce the bar chart of the BUSCO summary results. The locations of the BUSCO matches in the full_table.tsv files are converted into bigBed format for display on the GEP UCSC Genome Browser [available under the “BUSCO (diptera_odb10)” evidence track].

The *D. melanogaster* proteins, transcripts, and coding exons sequences were obtained from FlyBase (FlyBase release FB2022_03; *D. melanogaster* annotation release 6.46) and aligned against the *D. kikkawai*, *D. takahashii*, *D. bipectinata*, and *D. ananassae* Hi-C genome assemblies using the methods described under the “Muller F size estimation” section above. The SPALN alignments of *D. melanogaster* proteins against the Hi-C genome assemblies include the locations where insertions or deletions (indels) will result in frame-shifts. The locations of these potential frame-shifts are available under the “Potential Frame Shifts” evidence track on the GEP UCSC Genome Browser.

RefSeq gene models from *D. kikkawai*, *D. takahashii*, *D. bipectinata*, and *D. ananassae* were aligned against the corresponding Hi-C genome assembly using BLAT with the following parameters q=rna -fine -minScore=20 -stepSize=5. The transcript alignments were analyzed by pslReps with the parameters -minCover=0.15 -minAli=0.98 -nearTop=0.001, and then filtered by pslCDnaFIlter with the parameters -minId=0.95 -minCover=0.15 -localNearBest=0.001 -minQSize=20 -ignoreIntrons -repsAsMatch -ignoreNs -bestOverlap.

De novo repeat libraries were constructed for each species using Earl Grey (Baril *et al*. 2022). Repeat landscape plots were generated using the *createRepeatLandscape.pl* script from the RepeatMasker package (Smit *et al*. 2013).

### Wolbachia BLAST searches

We used the *wAna* genome (accession number: GCF_008033215.1) and for NCBI BLAST (Camacho *et al*. 2009), an E-value threshold of 1e-10 and a minimum alignment length of 1000 bp for *D. ananassae*. For *D. bipectinata*, we used an E-value threshold of 1e-5 and a minimum alignment length of 500 bp.

## Results and Discussion

### Long-read sequencing combined with Hi-C scaffolding results in chromosome-level scaffolds

We generated 12.9 to 13.7 Gb of Pacific Biosciences CLR reads for each species, which amounts to approximately 65x sequencing coverage based on a genome size of 205 Mb for each species [inferred from flow cytometry; (Gregory and Johnston 2008)] (Figure 2A). We compared the distributions of read lengths between species and calculated both the median and N50 values. For *D. bipectinata*, we obtained a median read length of 10.4 Kb and an N50 of 20.6 Kb. For *D. kikkawai*, the median read length was 13.1 Kb and read N50 was 32.1 Kb. For *D. takahashii*, the median read length was 10.3 Kb and read N50 was 20.7 Kb (Figures 2B and 2C). Read quality was assessed using SequelTools and SEQUELstats (see Methods, Supplementary Tables 2 and 3).

**Figure 2.**
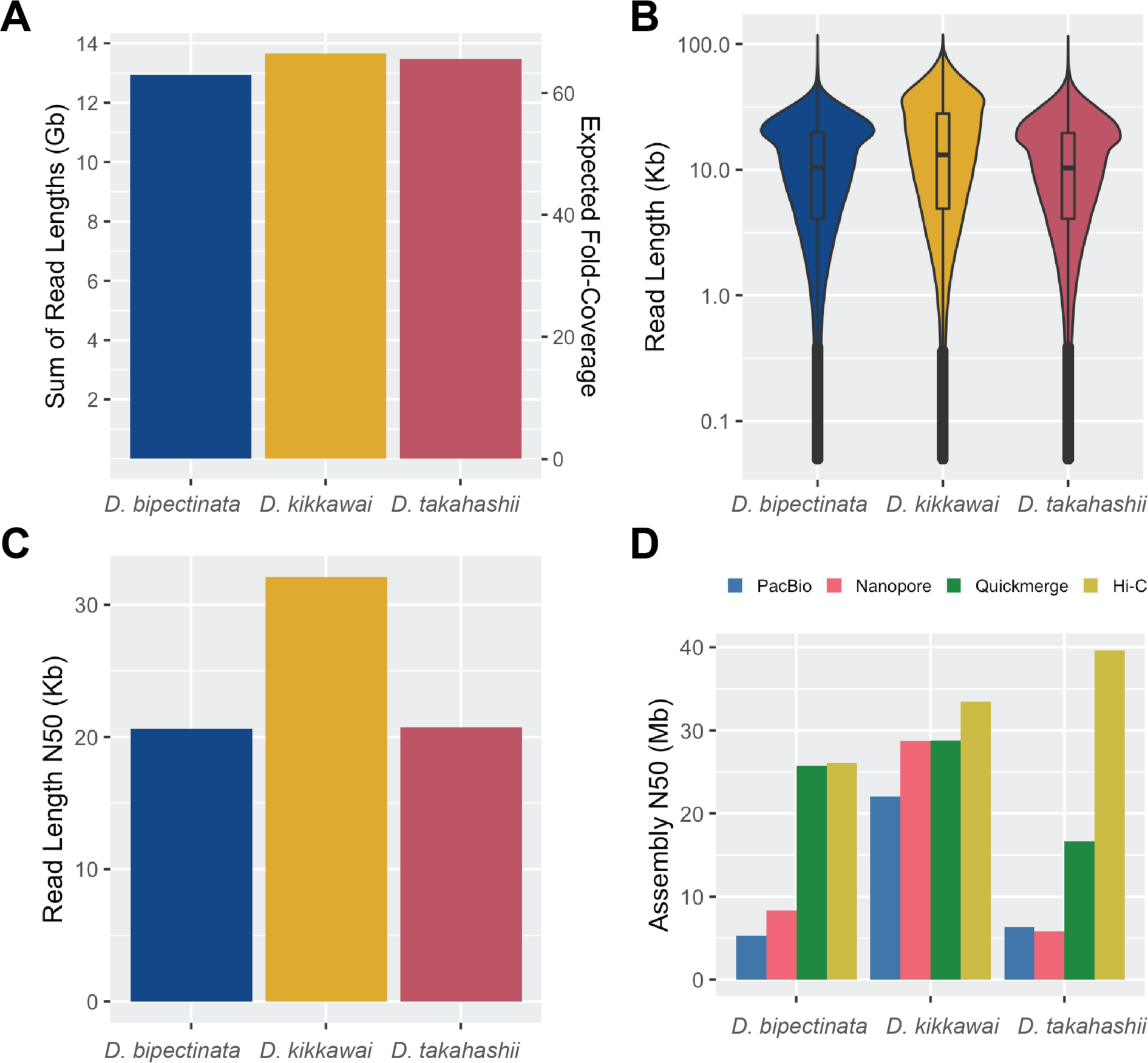
Summary of sequencing read lengths and assembly contiguity. (A) Total basepairs of Pacific Biosciences long read sequencing data generated for each species along with the expected sequencing coverage based on a genome size of 205 Mb for each species (estimated from flow cytometry) (B) Distribution of Pacific Biosciences read lengths for each species. (C) The read length N50 for each species. Half of the total sequencing data is present in reads of length N50 or larger. (D) Assembly N50 metrics for four different stages of the assembly pipeline: (1, blue) PacBio-only versions of each assembly were generated using Canu; (2, pink) Oxford Nanopore-only versions of each assembly were generated using Flye; (3, green) PacBio and Nanopore assemblies were merged with quickmerge; (4, yellow) the merged assemblies were scaffolded using Hi-C data.

**Figure 3.**
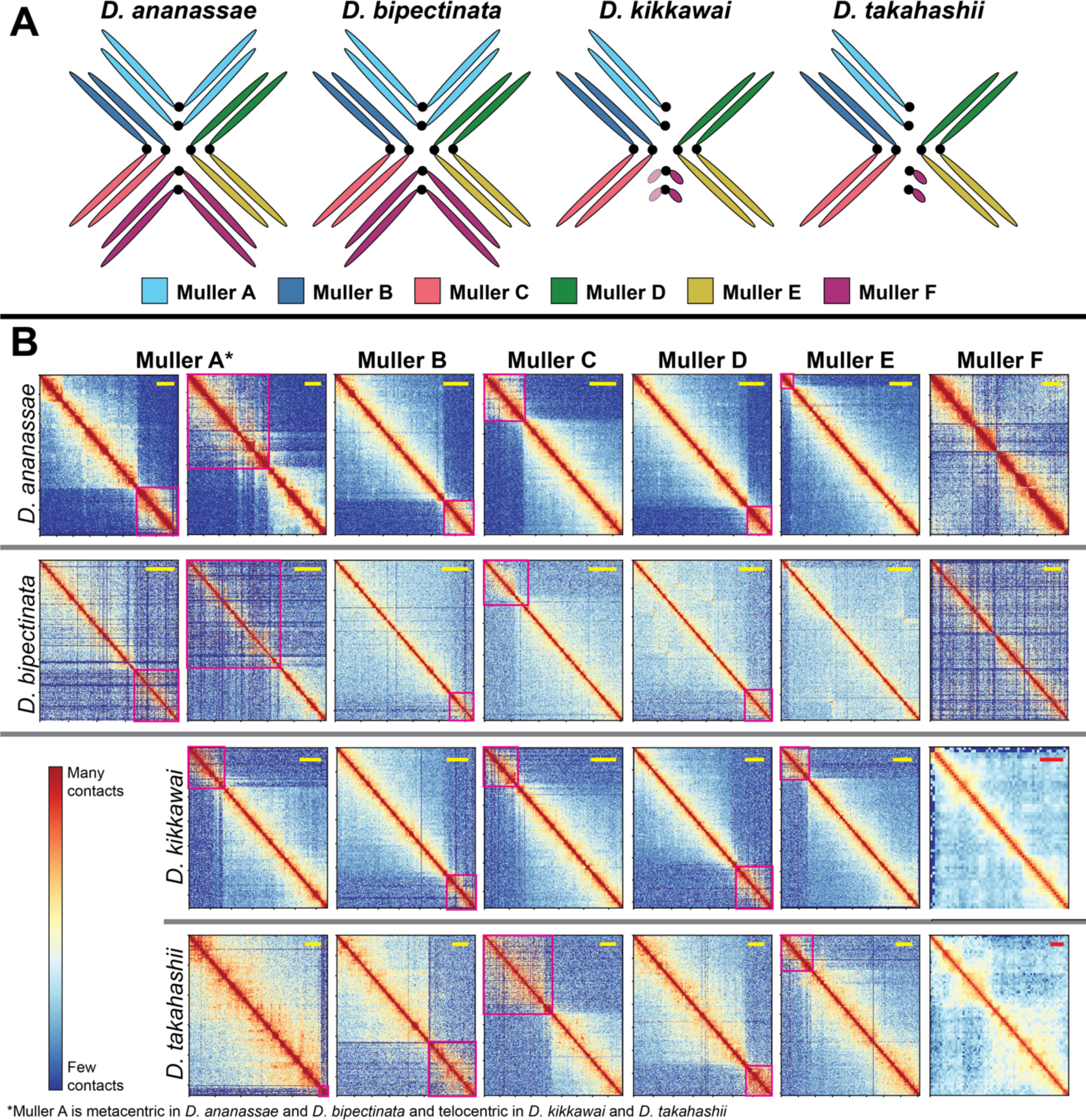
Muller Element Hi-C contact maps. (A) The Muller Elements A - F correspond to different chromosomes (or chromosome arms) in each species. Muller A is the X chromosome, which is telocentric in *D. melanogaster* but has become metacentric in *D. ananassae* and *D. bipectinata*. The B and C Elements are orthologous to the left and right arms of *D. melanogaster* chromosome 2, respectively. The D and E Elements are orthologous to the left and right arms of *D. melanogaster* chromosome 3, respectively. Published cytological data shows that the larger F elements in *D. ananassae* and *D. bipectinata* are metacentric, while the smaller F element in *D. takahashii* is telocentric, similar to *D. melanogaster* (Deng *et al*. 2007). Previous cytological studies have reported metacentric and telocentric F Elements in different populations of *D. kikkawai* (Baimai and Chumchong 1980). The *D. kikkawai* chromosome arm contains 93% (74/80) of the *D. melanogaster* F Element genes, and appears telocentric in our assembly. However, it is possible that the *D. kikkawai* F Element is actually metacentric but we were unable to assemble the other chromosome arm due to high repeat density. Note that chromosomes are not drawn to scale in this figure. (B) Hi-C contact maps are shown for each Muller Element (columns), for each species (rows). Pink boxes show the pericentromeric heterochromatin of each chromosome arm, which is spatially segregated from the euchromatin in the nucleus. The horizontal bars in the upper right corner of each panel are shown for scale: yellow bars represent 5 Mb while red bars represent 400 Kb. Note that two panels are shown for Muller A in *D. ananassae* and *D. bipectinata* because the chromosome is metacentric in these two species. Only one panel is shown for Muller A in *D. kikkawai* and *D. takahashii* because the chromosome is telocentric in these two species.

Previous work has shown assembly contiguity can be significantly improved by merging assemblies generated from different sequencing technologies and/or assembly algorithms (Alhakami *et al*. 2017). To implement this strategy, we generated two assemblies for each species using two different long-read sequencing platforms and assemblers: (1) contig assemblies for each species were generated using the PacBio reads and the *Canu* assembler (Koren *et al*. 2017), which resulted in contig N50 values of 5.3 Mb, 22.0 Mb, and 6.3 Mb for *D. bipectinata*, *D. kikkawai*, and *D. takahashii*, respectively (Figure 2D), and (2) contig assemblies were generated from Oxford Nanopore data [produced by Kim and colleagues (Kim *et al*. 2021)] using Flye, which resulted in contig N50 values of 8.4 Mb, 28.7 Mb, and 5.8 Mb for *D. bipectinata*, *D. kikkawai*, and *D. takahashii*, respectively. We then used *quickmerge* (Chakraborty *et al*. 2016) to merge our PacBio assemblies produced by Canu with our Nanopore assemblies produced by Flye. The merged assemblies showed improved contiguity with N50 values of 25.7 Mb, 28.8 Mb, and 16.6 Mb for *D. bipectinata*, *D. kikkawai*, and *D. takahashii*, respectively (Figure 2D).

We next used Hi-C data to scaffold our contig assemblies with the 3D-DNA pipeline (Dudchenko *et al*. 2017), which resulted in chromosome-level scaffolds for each species (Figure 3, Supplementary Table S4). We also generated Hi-C data for *D. ananassae*, which has an expanded F Element similar in size to that of *D. bipectinata*. We used the *D. ananassae* Hi-C data to scaffold the contigs from a recently published *D. ananassae* long-read genome assembly (Tvedte *et al*. 2021). Note that our use of “chromosome-level” nomenclature is based on the NCBI Assembly Data Model designation (see Methods).

To assess assembly completeness, we used BUSCO (Manni *et al*. 2021) to search for the presence of 3,285 dipteran single-copy orthologs. More than 99% of the single-copy orthologs were found in these assemblies, consistent with a high level of completeness (Figure 4). The *D. bipectinata* DbipHiC1 assembly has the highest percentage (1.2%) of “Complete (C) and duplicated (D)” genes among the four species. Examination of the duplicated BUSCO matches in the DbipHiC1 assembly shows that 27% (29/108) of the duplicated BUSCO matches correspond to Histone 2A (69968at7147) genes located at scaffold_165 and scaffold_175. Both scaffolds contain multiple copies of the histone genes His1, His2A, His2B, His3, and His4, which suggests that the duplicated genes in these scaffolds can be attributed to the histone gene cluster (Figure S1). Although these scaffolds were not detected by the *purge haplotigs* software package (see Methods), it remains possible that they represent two different haplotypes from the same locus (i.e. haplotigs), rather than separate histone arrays.

**Figure 4.**
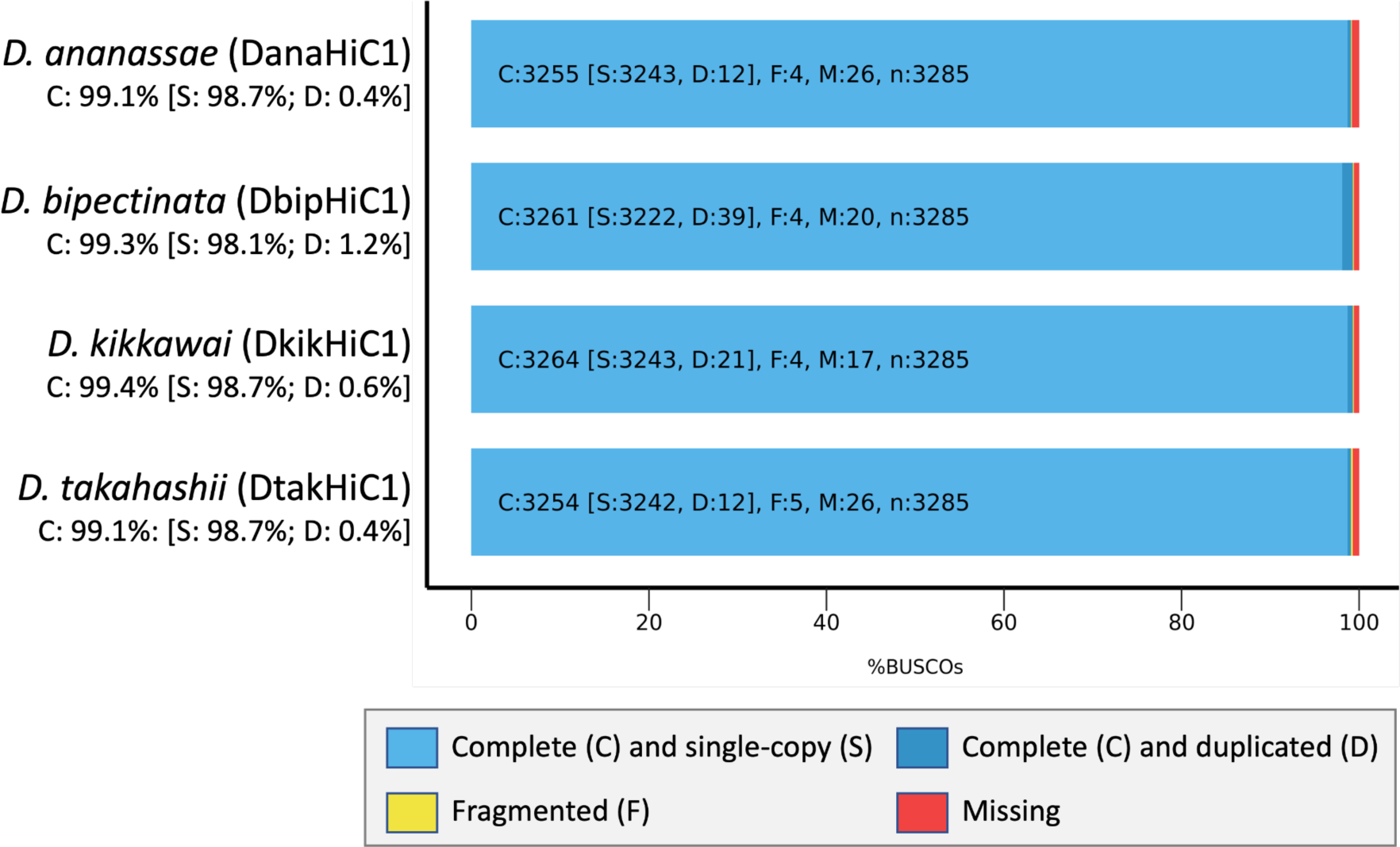
BUSCO short summary results. BUSCO analysis shows that 99.1% – 99.4% of the 3,285 single-copy orthologs in the diptera_odb10 lineage dataset are classified as “Complete” in the four Hi-C scaffolded genome assemblies. The percentages of Fragmented (F) and Missing (M) single-copy orthologs for each assembly are as follows: DanaHiC1 (F: 0.1%, M: 0.8%), DbipHiC1 (F: 0.1%, M: 0.6%), DkikHiC1 (F: 0.1%, M: 0.5%), DtakHiC1 (F: 0.2%, M: 0.8%).

In total, this work has resulted in chromosome-level genome assemblies for four *Drosophila* species with expanded F Elements (Figure S2).

### Chromosome size variation among species

The chromosome-level scaffolds produced by this study allow us to compare the sizes of chromosome arms, including pericentromeric heterochromatin, among species (Figure 5). The total assembly sizes for each species are 192.2 Mb (*D. ananassae*), 194.5 Mb (*D. bipectinata*), 188.5 Mb (*D. kikkawai*), and 198.1 Mb (*D. takahashii*), all close to, but slightly less than, the 205 Mb size estimate from flow cytometry (Gregory and Johnston 2008). The size of the F Element scaffold in each species is 19.4 Mb (*D. ananassae*), 20.5 Mb (*D. bipectinata*), 2.3 Mb (*D. kikkawai*), and 3.2 Mb (*D. takahashii*). F Element expansion (compared to *D. melanogaster*) in these species therefore ranges from 1.8-fold (*D. kikkawai*) to 15.8-fold (*D. bipectinata*). Interestingly, despite the large increase in size of the F Element in *D. ananassae* and *D. bipectinata*, the estimated total genome sizes based on the chromosome-level assemblies are similar across all four species (Figure 5). In fact, the assembled portions of the Muller Elements B, C, D and E are all smaller in both *D. ananassae* and *D. bipectinata* compared to *D. kikkawai* and *D. takahashii* (Figure 5). Note, however, that these genome size estimates are derived from the sizes of the assembled scaffolds, and they could be confounded by differences in the number of sequences that cannot be assembled or scaffolded due to high repeat content in the four *Drosophila* species. Thus, in all cases these are minimal estimates of Muller Element size.

**Figure 5.**
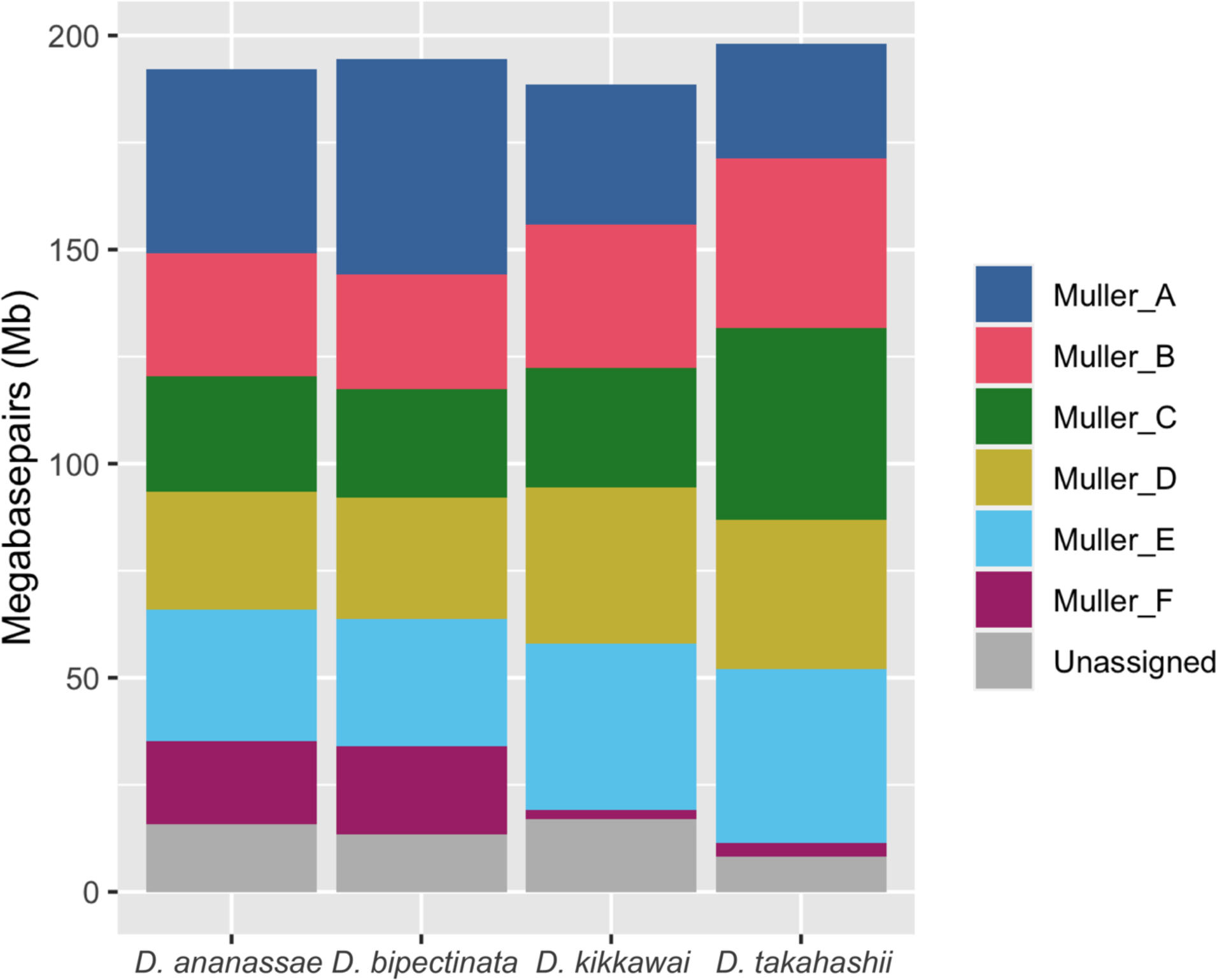
Scaffold sizes for each species. The size in megabases is shown for each Muller Element scaffold in the Hi-C assemblies for the four *Drosophila* species. Scaffolds that were not assigned to Muller Elements are grouped together in the “Unassigned” category. For each species, the combined height of the colored boxes is equal to the assembly size.

### Annotation of genes and repetitive elements

Students participating in the Genomics Education Partnership (GEP) will manually construct gene models for the F Element and for a region near the base of the D Element for the four *Drosophila* species discussed here using RNA-Seq data, computational gene predictions, and sequence similarity to *D. melanogaster* genes.

The high error rate of PacBio continuous long-read (CLR) sequencing data can lead to consensus errors in the resulting assembly. When found within gene coding sequences, these consensus errors can cause artifactual frameshift mutations, which decrease the accuracy of computational gene predictions. As part of the assessment of the quality of the Hi-C scaffolded genome assemblies, 30,799 proteins from 13,986 genes in *D. melanogaster* annotation release 6.46 provided by FlyBase (Gramates *et al*. 2022) were aligned against each genome assembly. Between 91% to 96% of the *D. melanogaster* isoforms have at least one SPALN alignment in the Hi-C scaffolded genome assemblies, accounting for 86% to 93% of the *D. melanogaster* genes (Table 1). The number of frameshifts in the SPALN protein alignments range from 507 to 552, representing ∼4% of genes. The number of frameshifts is similar across all four assemblies, despite the fact that the *D. ananassae* contigs were generated from high-accuracy PacBio HiFi sequencing data (Tvedte et al. 2021). This comparison suggests that our *D. bipectinata*, *D. kikkawai*, and *D. takahashii* assemblies do not suffer from a high rate of consensus errors, despite being generated from lower accuracy PacBio CLR data.

**Table 1.** SPALN alignments show that 86–93% of the protein-coding genes from *D. melanogaster* annotated release 6.46 (which consists of 30,799 proteins derived from 13,986 genes) can be placed in the Hi-C scaffolded genome assemblies.

| Species (Assembly) | Number of Genes with Alignments | Number of Isoforms with Alignments | Estimated Number of Frame Shifts |
| --- | --- | --- | --- |
| *D. kikkawai* (DkikHiC1) | 12,059 (86%) | 27,997 (91%) | 507 |
| *D. takahashii* (DtakHiC1) | 12,045 (86%) | 27,970 (91%) | 503 |
| *D. bipectinata* (DbipHiC1) | 12,119 (87%) | 28,097 (91%) | 512 |
| *D. ananassae* (DanaHiC1) | 13,037 (93%) | 29,456 (96%) | 552 |

In the *D. melanogaster* annotation release 6.46, the F Element has 298 isoforms derived from 80 genes. Between 93% to 98% of the *D. melanogaster* F Element isoforms have at least one SPALN alignment in the Hi-C scaffolded genome assemblies, accounting for the 91% to 96% of the F Element genes (Table 2). The number of frame shifts in the protein alignments to F Element genes ranges from 1 to 5.

**Table 2.** SPALN alignments shows that 91–96% of the F Element protein-coding genes from *D. melanogaster* annotated release 6.46 (which consists of 298 proteins derived from 80 genes) can be placed in the Hi-C scaffolded genome assemblies.

| Species (Assembly) | Number of Genes with Alignments | Number of Isoforms with Alignments | Estimated Number of Frame Shifts |
| --- | --- | --- | --- |
| *D. kikkawai* (DkikHiC1) | 74 (93%) | 287 (96%) | 5 |
| *D. takahashii* (DtakHiC1) | 77 (96%) | 293 (98%) | 1 |
| *D. bipectinata* (DbipHiC1) | 73 (91%) | 281 (94%) | 1 |
| *D. ananassae* (DanaHiC1) | 73 (91%) | 278 (93%) | 4 |

We used Earl Grey (Baril *et al*. 2022) to create *de novo* repeat libraries for each species. The number of families identified and sum of consensus lengths for each species are as follows: in *D. ananassae*, 1118 families sum to 4.5 Mb, in *D. bipectinata*, 1014 families sum to 3.8 Mb, in *D. kikkawai*, 924 families sum to 3.5 Mb, and in *D. takahashii*, 1066 families sum to 3.5 Mb. We then used RepeatMasker (Smit *et al*. 2013) to identify the locations of each repeat family within their respective genome assemblies. Using our custom repeat libraries, RepeatMasker masked a total of 88.8 Mb (41.5%) of the *D. ananassae* genome assembly, 69.3 Mb (35.6%) of the *D. bipectinata* genome assembly, 58.6 Mb (31.1%) of the *D. kikkawai* genome assembly, and 60.3 Mb (30.4%) of the *D. takahashii* genome assembly. We then used the sequence divergence among individual insertions from the same repeat family, along with the percentage of the genome occupied by each family, to visualize the repeat landscape for each species (Figure 6).

**Figure 6.**
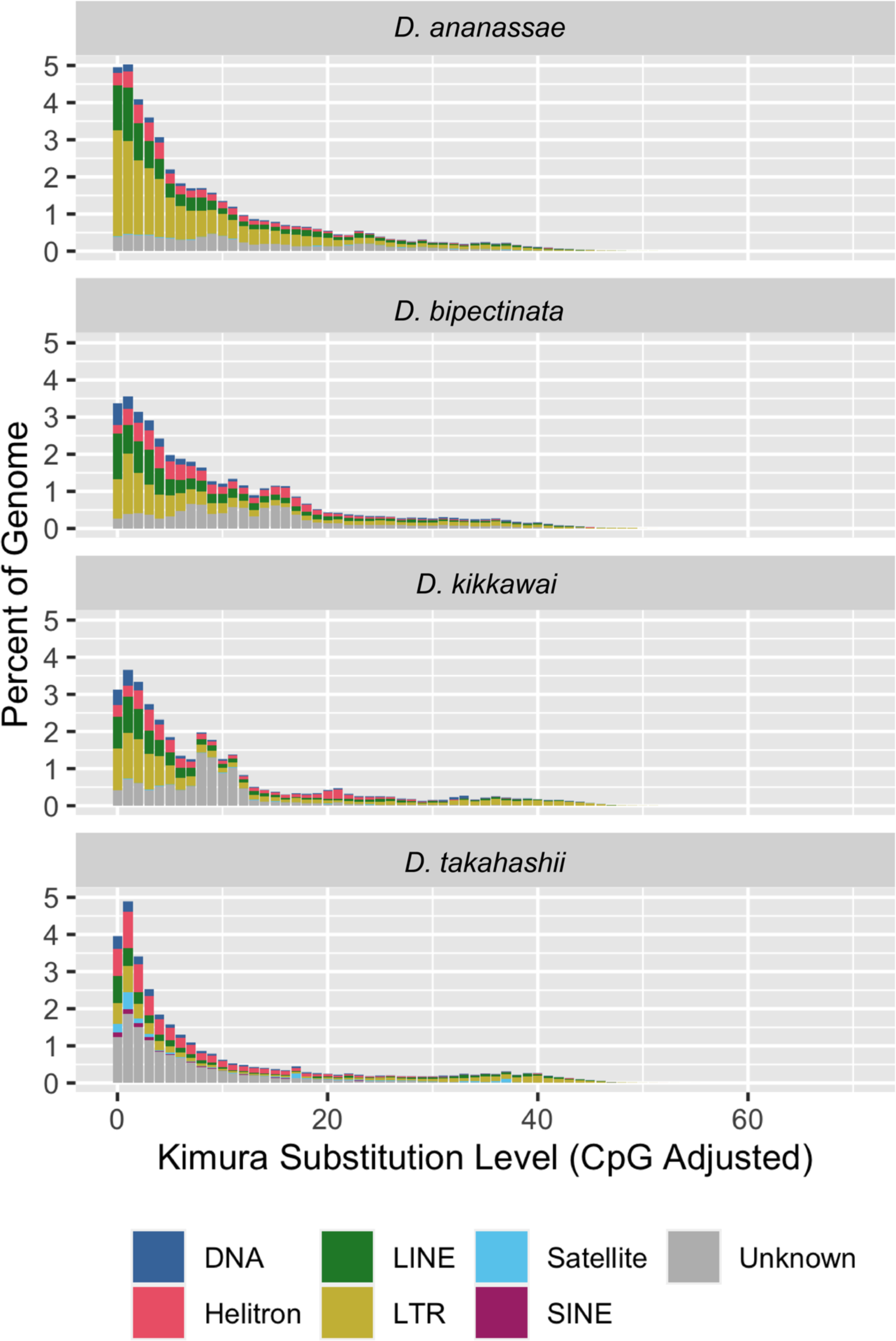
Repeat landscape plots. Repeat landscapes were generated from *de novo* repeat libraries created for each species using the assemblies reported here. The x-axis shows the sequence divergence among individual copies from the same repeat family, corrected using the Kimura 2 Parameter model. Each bar summarizes the percent of the genome occupied by each repeat superfamily/subclass for a given divergence level.

### Contribution of *Wolbachia* to F Element expansion

Previous work has suggested that lateral transfer of DNA from the *Wolbachia* endosymbiont into the nuclear genome of *D. ananassae* has contributed to the size expansion of the F Element in this species (Klasson *et al*. 2014). A recent study constructed a long-read genome assembly (accession number: GCF_017639315.1) from a strain of *D. ananassae* that was treated with tetracycline and cured of *Wolbachia* infections. The study showed that approximately 4.9 Mb of Wolbachia sequences have integrated into the *D. ananassae* genome (Tvedte *et al*. 2022), likely in the F Element (Tvedte et al. 2021).

Our Hi-C data allowed us to place many contigs within chromosome-level scaffolds. We therefore sought to determine whether the Hi-C scaffolding process allowed the *Wolbachia* sequences previously identified in the *D. ananassae* assembly to be assigned to one or more Muller Element scaffolds. We also investigated whether *Wolbachia* sequence was present within our *D. bipectinata* assembly, which shows a level of F Element expansion similar to that of *D. ananassae*. We used BLAST (Camacho *et al*. 2009) to search the complete *wAna Wolbachia* genome assembly (accession number: GCF_008033215.1) against the Hi-C scaffolded *D. ananassae* assembly reported here. This BLAST search identified a similar amount of *Wolbachia* sequence as the previous study (4.97 Mb). However, all of the significant BLAST hits to *wAna* were on scaffolds that could not be assigned to *D. ananassae* Muller Elements using the Hi-C data; in particular, there were no significant BLAST hits to the *D. ananassae* F Element scaffold.

We also performed a BLAST search of the *wAna* genome assembly against the Hi-C scaffolded *D. bipectinata* assembly, using less stringent parameters to account for the possibility that the *D. bipectinata* nuclear genome contains DNA from a different *Wolbachia* subtype. In contrast to the *D. ananassae* results, we only identify ∼19 Kb of sequence in the *D. bipectinata* assembly that matches the *wAna* genome. All of the matches are located in scaffolds that could not be assigned to the *D. bipectinata* Muller Elements.

Collectively, our analysis of the Hi-C scaffolded assemblies cannot rule out potential contributions of horizontal transfer of *Wolbachia* DNA to the expansion of the *D. ananassae* F Element. However, our results strongly suggest that horizontal transfer of *Wolbachia* DNA is not a major contributor to the expansion of the *D. bipectinata* F Element. Furthermore, incorporation of *Wolbachia* DNA would explain, at most, ∼20% of the expansion of the Muller F Element in *D. ananassae*, which means that ∼80% of the size increase is due to other factors, such as accumulation of mobile DNA and other repeats.

## Data Availability Statement

The PacBio sequencing data are available through the NCBI BioProject database under the accession number PRJNA948012 and the Hi-C data are available through accession numbers PRJNA961071 and PRJNA967347.

The Hi-C scaffolded genome assemblies for *D. bipectinata*, *D. takahashii*, *D. kikkawai*, and *D. ananassae* have been deposited at GenBank under the accession numbers JARPSB000000000, JARPSC000000000, JARPSD000000000, and JASIRA000000000, respectively. The versions of the *D. bipectinata*, *D. takahashii*, *D. kikkawai*, and *D. ananassae* genome assemblies described in this paper are JARPSB010000000, JARPSC010000000, JARPSD010000000, and JASIRA010000000 respectively.

The genome assemblies and evidence tracks described in this manuscript are displayed on the GEP UCSC Genome Browser, available through the links under the “UCSC Genome Browser” column of the “<u>Hi-C Genome Assemblies for F Element Expansion Project</u>” landing page. These Genome Browsers include additional gene predictions, repeat analysis, and RNA-Seq evidence tracks that will be used in the subsequent comparative analyses of the expansion of the F Elements.

## Acknowledgements

The authors would like to thank Professor Artyom Kopp (UC Davis) for providing the flies used for the PacBio sequencing and for clarifying the history of the strains of *D. kikkawai*, *D. takahashii*, and *D. bipectinata* used by the different sequencing projects. We would also like to thank Dr. Bernard Kim (Stanford University) for providing the high molecular weight DNA used for the PacBio sequencing, and the McDonnell Genome Institute (MGI) for performing the PacBio sequencing. The authors acknowledge the Office of Advanced Research Computing (OARC) at Rutgers, The State University of New Jersey for providing access to the Amarel cluster and associated research computing resources that have contributed to the results reported here.

## Supplemental Materials

**Figure S1.**
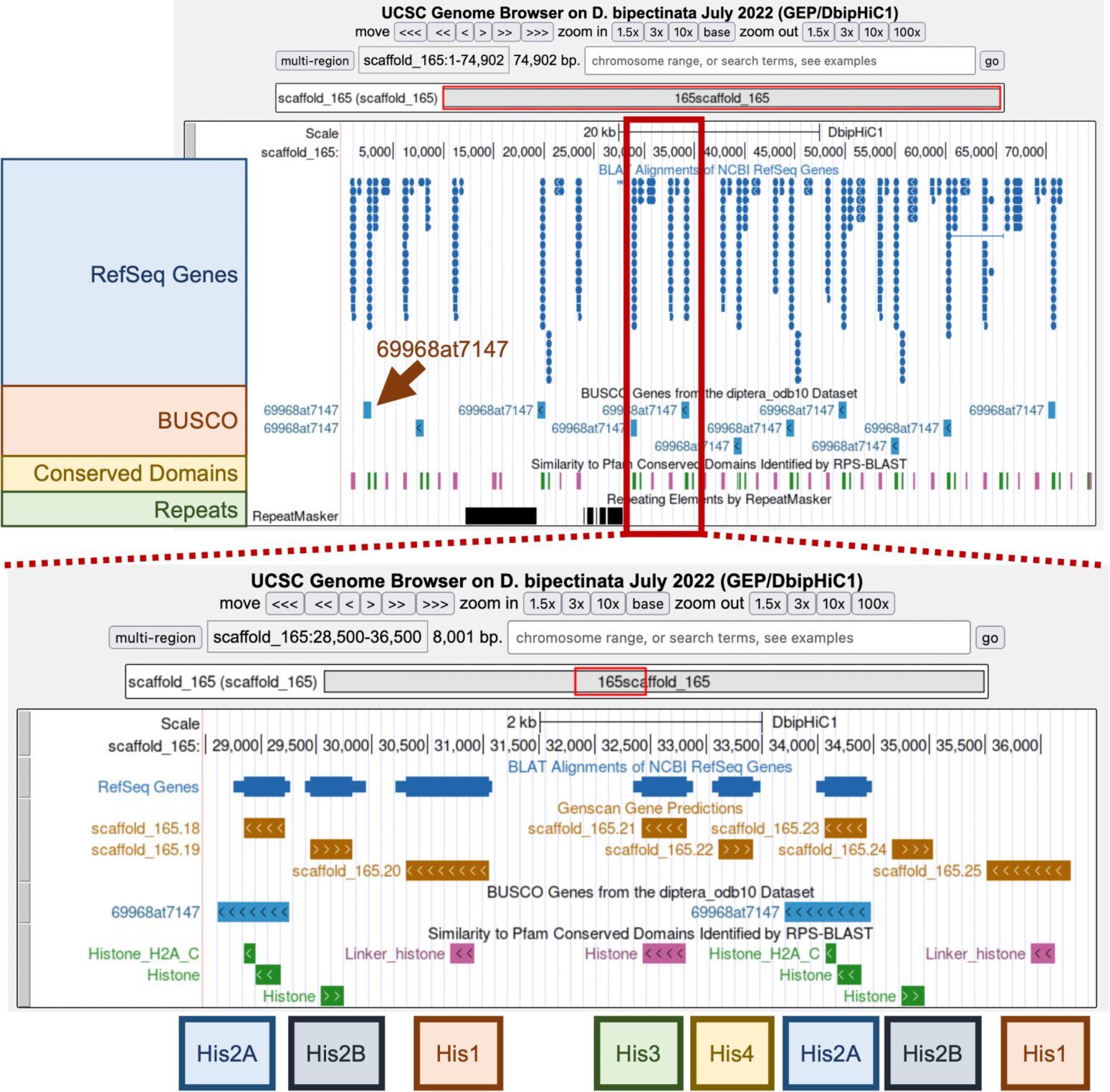
Complete and duplicated BUSCO matches in the *D. bipectinata* Hi-C assembly can partly be attributed to the histone gene cluster on scaffold_165 and scaffold_175. Among the 108 complete and duplicated BUSCO matches in the *D. bipectinata* Hi-C assembly, 29 of them (27%) are located on scaffold_165 and scaffold_175. (Top) Examination of the entire 75 Kb scaffold_165 shows multiple matches to Histone 2A (69968at7147 in the diptera_odb10 lineage dataset; brown arrow). (Bottom) Examination of the 28,500-36,500 region of scaffold_165 shows the histone gene cluster with copies of the genes for Histones 3 (*His3*), 4 (*His4*), 2A (*His2A*), 2B (*His2B*), and 1 (*His1*).

**Figure S2.**
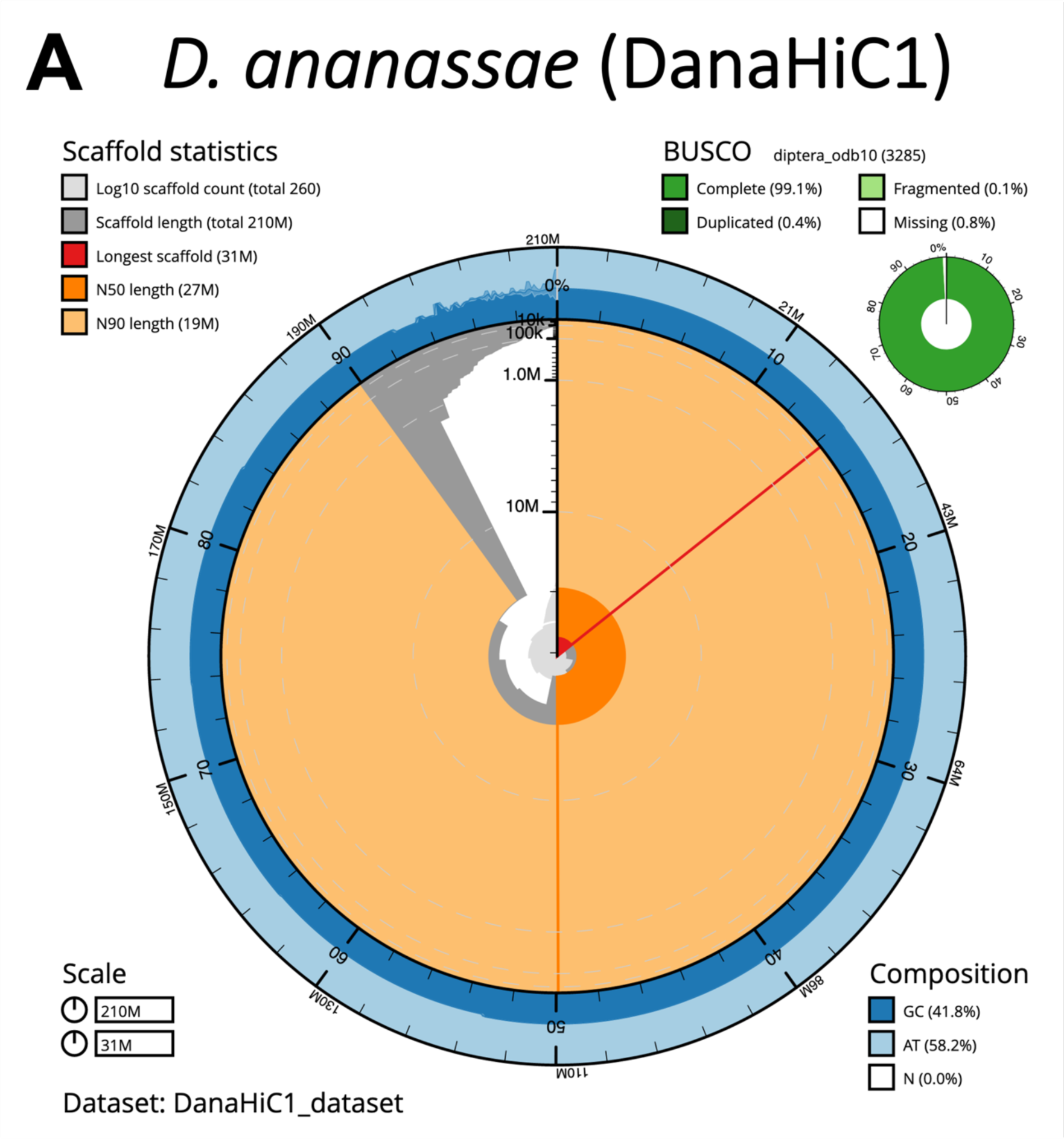
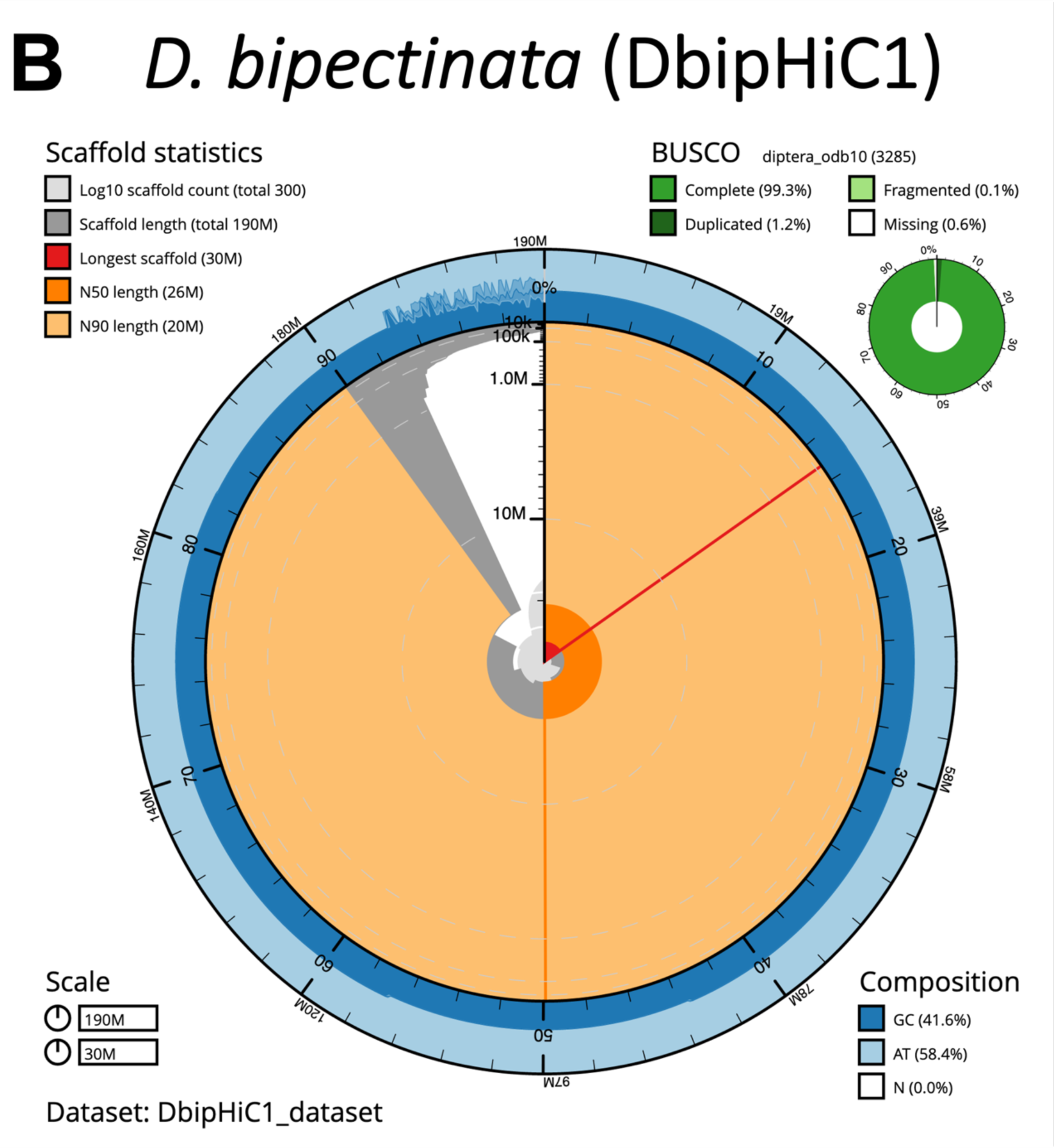
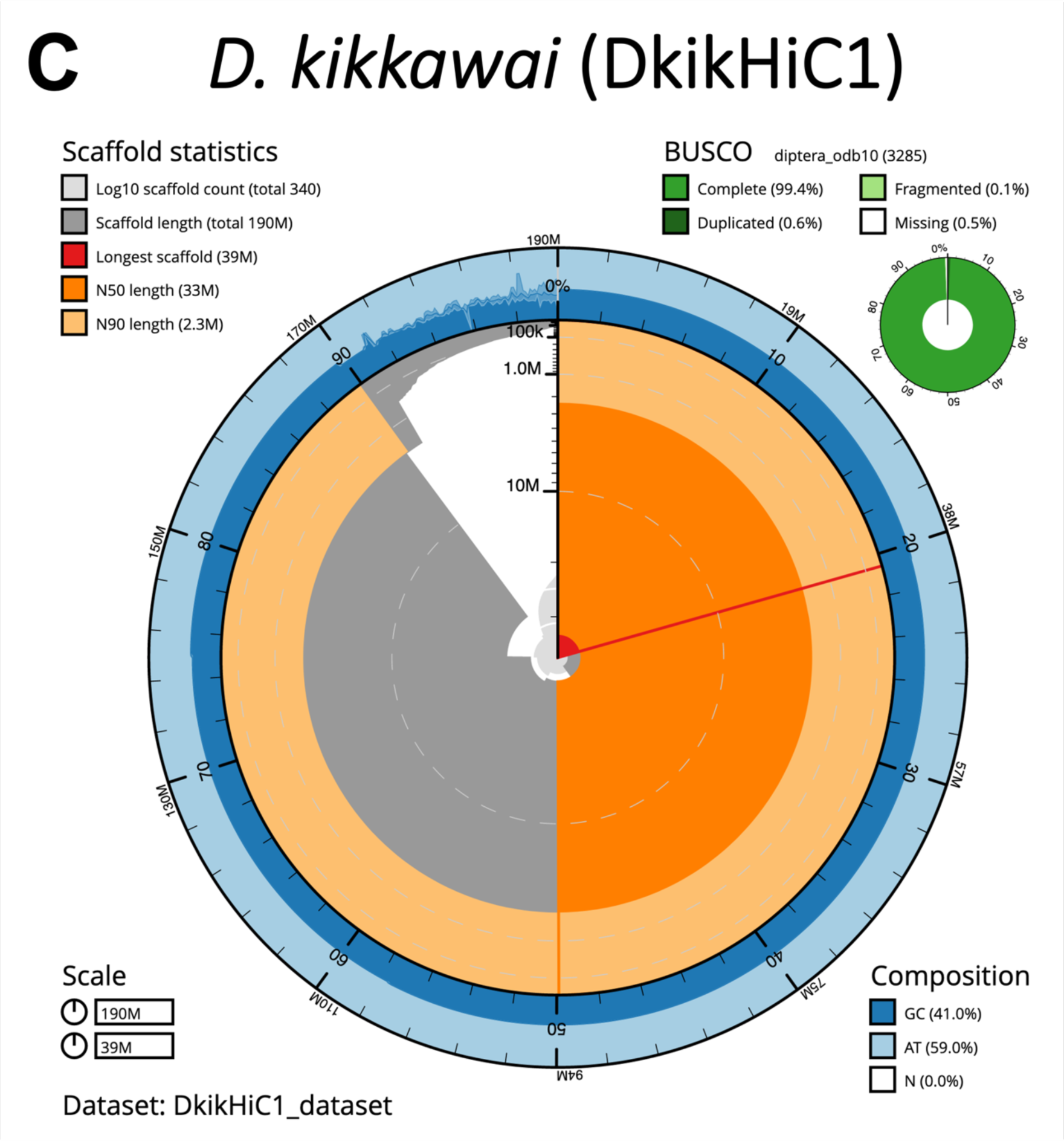
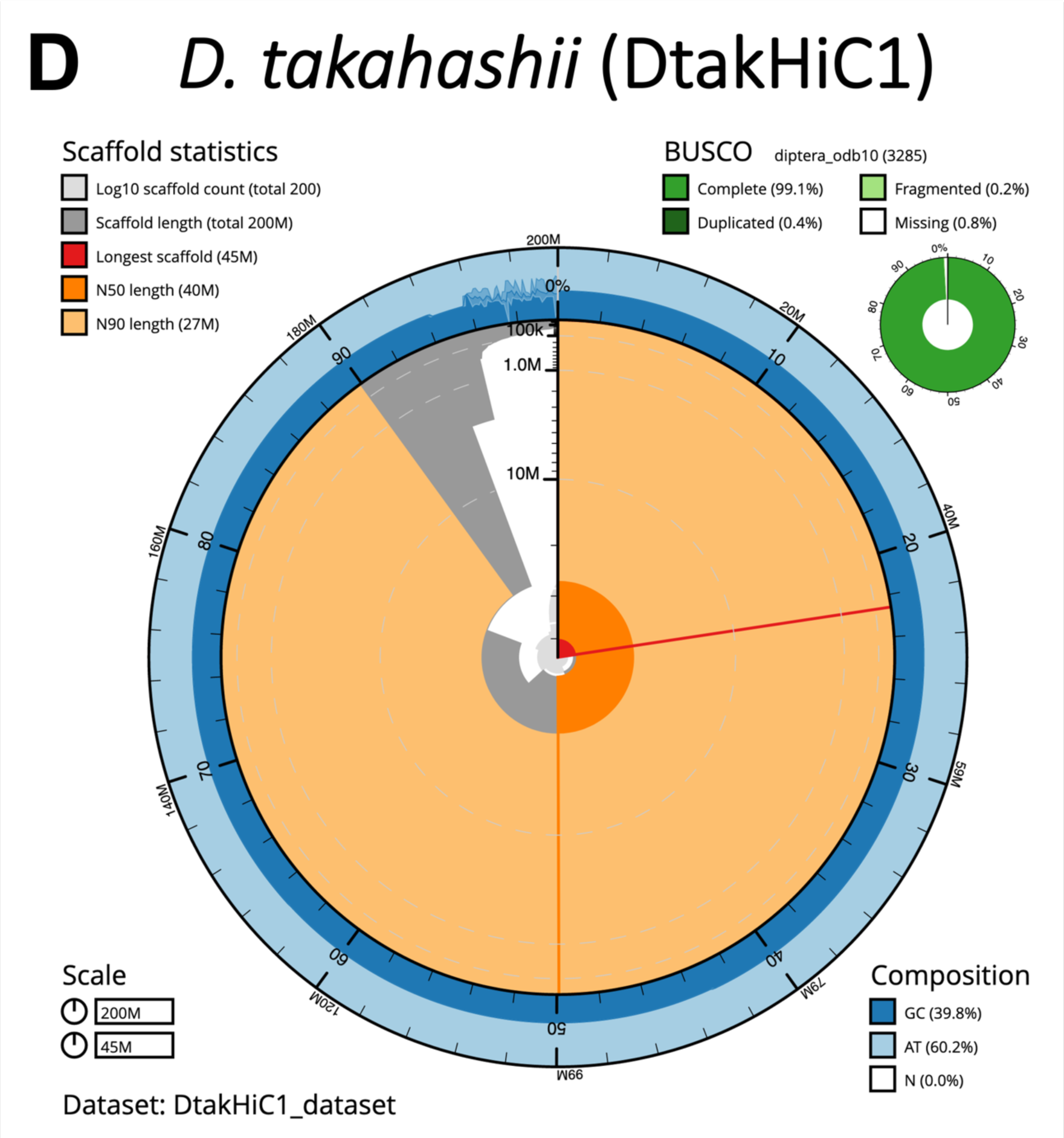
Summary of Hi-C scaffolded assemblies. Snail plots which show the assembly statistics for *D. ananassae* (A), *D. bipectinata* (B), *D. kikkawai* (C), and *D. takahashii* (D). The Hi-C scaffolded assemblies for the four genomes are high quality — with scaffold N50s that range from 26 Mb in *D. bipectinata* to 40 Mb in *D. takahashii* and the percentages of “Complete” BUSCOs range from 99.1% in *D. takahashii* and *D. ananassae* to 99.4% in *D. kikkawai*.

**Table S1.** Version information and references for the Bioinformatics tools used in this study.

| Tool | Version | URL | Reference |
| --- | --- | --- | --- |
| 3D-DNA | 180419 | https://github.com/aidenlab/3d-dna | PMID: 28336562 |
| BBMap | 38.86 | https://sourceforge.net/projects/bbmap/ |  |
| BlobTools2 | 3.2.7 | https://blobtoolkit.genomehubs.org/ | PMID: 32071071 |
| BUSCO | 4.1.4 | https://gitlab.com/ezlab/busco | PMID: 34320186 |
| Canu | 2.1 | https://github.com/marbl/canu | PMID: 28298431 |
| Earl Grey | Git commit<br>2dcdb8b | https://github.com/TobyBaril/EarlGrey | DOI: 10.21203/rs.3.rs-1812599/v1 |
| Flye | 2.8.1 | https://github.com/fenderglass/Flye | PMID: 30936562 |
| GCpp | 1.0.0-1807624 | https://github.com/PacificBiosciences/gcpp |  |
| GEP UCSC<br>Genome Browser | 435 | https://gander.wustl.edu |  |
| Hapo-G | Git commit<br>ca2c732 | https://github.com/institut-de-genomique/HAPO-G | PMID: 33987534 |
| HiCEXplorer | 3.6 | https://github.com/deeptools/HiCEXplorer | PMID: 29335486 |
| LAST | 1406 | https://gitlab.com/mcfrith/last | PMID: 21209072 |
| NCBI BLAST+ | 2.13.0 | https://blast.ncbi.nlm.nih.gov/Blast.cgi | PMID: 20003500 |
| NextPolish | 1.3.0 | https://github.com/Nextomics/NextPolish | PMID: 31778144 |
| pbmm2 | 1.1.0 | https://github.com/PacificBiosciences/pbmm2 |  |
| POLCA | 3.4.2 | https://github.com/alekseyzimin/masurca | PMID: 32589667 |
| Porechop | 0.2.4 | https://github.com/rrwick/Porechop |  |
| Purge Haplotigs | 1.1.1 | https://bitbucket.org/mroachawri/purge_haplotigs/ | PMID: 30497373 |
| quickmerge | 0.3 | https://github.com/mahulchak/quickmerge | PMID: 27458204 |
| RepeatMasker | 4.1.2-p1 | http://repeatmasker.org/ |  |
| SEQUELstats | Git commit<br>452194f | https://github.com/VertebrateResequencing/SEQUELstats |  |
| SequelTools | Git commit<br>553b9fe | https://github.com/ISUgenomics/SequelTools | PMID: 33004007 |
| SMRT Link | 7.0.1.66975 | https://www.pacb.com/support/software-downloads/ |  |
| SPALN | 2.3.3f | https://github.com/ogotoh/spaln | PMID: 22848105 |
| UCSC Genome<br>Browser | 435 | https://hgdownload.soe.ucsc.edu/admin/exe/ | PMID: 36420891 |
| WU-BLAST | 2.0 | https://blast.advbiocomp.com/ |  |

